# Somatic CAG expansion in Huntington’s disease is dependent on the MLH3 endonuclease domain, which can be excluded via *MLH3* splice redirection to suppress expansion

**DOI:** 10.1101/2020.10.26.356238

**Authors:** Jennie C. L. Roy, Antonia Vitalo, Marissa A. Andrew, Eduarda Mota-Silva, Marina Kovalenko, Zoe Burch, Anh M. Nhu, Paula E. Cohen, Ed Grabczyk, Vanessa C. Wheeler, Ricardo Mouro Pinto

## Abstract

Somatic expansion of the CAG repeat tract that causes Huntington’s disease (HD) is thought to contribute to the rate of disease pathogenesis. Therefore, factors influencing repeat expansion are potential therapeutic targets. Genes in the DNA mismatch repair pathway are critical drivers of somatic expansion in HD mouse models. Here, we have tested, using genetic and pharmacological approaches, the role of the endonuclease domain of the mismatch repair protein MLH3 in somatic CAG expansion in HD mice and patient cells. A point mutation in the MLH3 endonuclease domain completely eliminated CAG expansion in the brain and peripheral tissues of a HD knock-in mouse model (*Htt*^Q111^). To test whether the MLH3 endonuclease could be manipulated pharmacologically, we delivered splice switching oligonucleotides in mice to redirect *Mlh3* splicing to exclude the endonuclease domain. Splice redirection to an isoform lacking the endonuclease domain was associated with reduced CAG expansion. Finally, CAG expansion in HD patient-derived primary fibroblasts was also significantly reduced by redirecting *MLH3* splicing to the endogenous endonuclease domain-lacking isoform. These data indicate the potential of targeting the MLH3 endonuclease domain to slow somatic CAG repeat expansion in HD, a therapeutic strategy that may be applicable across multiple repeat expansion disorders.

## Introduction

Huntington’s disease (HD) is a dominantly inherited neurodegenerative disorder caused by the inheritance of an expanded CAG trinucleotide repeat in the huntingtin gene (*HTT*), encoding an extended polyglutamine tract in the huntingtin protein (1). The expanded CAG mutation ultimately results in neuronal dysfunction and death (2), via mechanisms that are as yet unclear (3). The mutant CAG repeat undergoes further expansion in somatic cells in a CAG length-dependent and tissue- or cell-type-specific manner (4–8). Somatic expansion is recapitulated in HD mouse models that also show progressive repeat tract lengthening over time (9–12), with the striatum and liver showing high levels of expansion, particularly the medium spiny neurons and hepatocytes, respectively (9, 10). Studies of postmortem HD brains revealed that longer somatic CAG expansions are associated with an earlier age of disease onset (7), with more recent studies showing correlations between age of onset or disease progression and the degree of somatic expansion in HD patient blood (13). Significantly, recent genome-wide association studies (GWAS) of over 9,000 HD patients have demonstrated that the rate of disease onset is determined by the length of the pure CAG repeat, rather than the length of the glutamine tract in huntingtin (14). This is supported by additional studies (13, 15) and provides compelling support for a model in which the rate of somatic CAG expansion drives the rate of disease onset. Thus, therapies targeting somatic repeat expansions have the potential to delay disease onset.

Genetic knockout studies in HD mouse models have shown that *Msh2, Msh3, Mlh1*, and *Mlh3* genes in the DNA mismatch repair (MMR) pathway are required for somatic expansion (16–19). Human GWAS have identified several genes in this pathway (*MSH3, MLH1, PMS1, PMS2*) to be associated with age at motor onset (14, 20), with *MSH3* also showing association with a measure of HD progression (21). The mouse ortholog of an additional HD onset-modifier gene, *FAN1* (14, 20), with no known function in the MMR pathway, suppressed somatic expansion in HD knock-in mice in a manner that was dependent on *Mlh1* (8). Genetic variation in some human onset or progression modifier genes (*MSH3, MLH1, MLH3, FAN1*) also modified CAG expansion in HD patient blood or blood-derived cells (13, 14, 22). Together, these data indicate that HD pathogenesis is modulated by naturally occurring variation in genes that act by altering the rate of somatic CAG expansion, and that genes driving this process are potential therapeutic targets for HD. MMR pathway genes also modify somatic expansion in mouse models and in mouse or human cell-based models of a number of other DNA repeat expansion disorders, including myotonic dystrophy type I (DM1) (23–25), Friedreich ataxia (FRDA) (26–29), and the fragile X-related disorders (FXD) (30–34). In addition, polymorphisms in two DNA repair genes (*FAN1, PMS2*) associated with HD onset were also associated with disease onset in the spinocerebellar ataxias (35). Thus, therapeutic strategies targeting MMR genes may be applicable to multiple repeat expansion diseases.

Our previous genetic knockout studies in *Htt*^Q111^ knock-in mice demonstrated that MLH3 is an essential component of the CAG repeat expansion process (17). MLH3 is known to play an important role in meiosis, but a relatively minor role in canonical MMR (36–38). MLH3 contains an evolutionary conserved endonuclease domain (39) that we postulated may be important in its CAG expansion-promoting role in HD (17), with inactivation or elimination of this domain hypothesized to eliminate or reduce somatic expansions. Interestingly, human MLH3 exists primarily as two protein isoforms: isoform 1 (UniProt Q9UHC1-1) is specified by splice variant 1 (CCDS32123) containing exon 7, which encodes the endonuclease domain, while isoform 2 (UniProt Q9UHC1-2) is specified by splice variant 2 (CCDS9837), lacking exon 7, and therefore has no endonuclease domain (28, 40). *MLH3* splice redirection from predominantly splice variant 1 to predominantly splice variant 2 can be achieved with splice switching oligonucleotides (SSOs) that mask the acceptor and donor sites that surround exon 7 of the *MLH3* pre-mRNA (28). In a previous FRDA study, splice redirection of *MLH3* to exclude the endonuclease domain resulted in reduced rates of GAA expansion in a human cell-based model and in patient cells (28). This indicated that the MLH3 endonuclease domain was important for GAA repeat expansion and that splice redirection could be used to reduce its contribution to the expansion process.

Here we have tested the role of the MLH3 endonuclease domain in somatic *HTT* CAG expansion, using both genetic studies and a pharmacological approach based on splice redirection. We demonstrate that: 1) a single point mutation in the MLH3 endonuclease domain, predicted to abrogate endonuclease activity, eliminates somatic CAG expansion in *Htt*^Q111^ mice; 2) systemic delivery of SSOs to redirect *Mlh3* splicing to exclude the endonuclease domain suppresses somatic CAG expansion in *Htt*^Q111^ mice; and 3) *MLH3* splice redirection with SSOs also effectively reduces CAG expansion in HD patient-derived primary fibroblast cells, in the absence of replication. These data highlight the therapeutic potential of targeting the MLH3 endonuclease domain to slow *HTT* CAG repeat expansion, a strategy that may be applicable across multiple repeat expansion disorders.

## Materials and Methods

### Mice

*Htt*^Q111^ HD knock-in mice (formerly *Hdh*^Q111^) (41) were maintained on a C57BL/6J background (9) by breeding heterozygous males to C57BL/6J wild-type females from The Jackson Laboratory (Bar Harbor, ME). *Mlh3^DN^* mice (42) were also maintained on C57BL/6J background. *Htt*^Q111/+^ and *Mlh3*^WT/DN^ mice were crossed together to generate *Htt*^Q111/+^*Mlh3*^WT/DN^ and *Htt*^+/+^ *Mlh3*^WT/DN^ mice, which were subsequently intercrossed to generate heterozygous *Htt*^Q111/+^ mice (hereafter referred to as *Htt*^Q111^, for simplicity) that were either wild type (*Mlh3*^WT/WT^), heterozygous (*Mlh3*^WT/DN^), or homozygous (*Mlh3*^DN/DN^) for the *Mlh3^DN^* point mutation. Genotyping of the *Htt*^Q111^ and *Mlh3*^DN^ alleles in genomic DNA extracted from tail or ear biopsies at weaning, or in adult tissues for genotype confirmation, was carried out as previously described (16, 42). All animal procedures were carried out to minimize pain and discomfort, under approved IACUC protocols of the Massachusetts General Hospital. Animal husbandry was performed under controlled temperature and light/dark cycles.

### *HTT* CAG repeat instability analysis

Genomic DNA was isolated from mouse tissues and patient fibroblast cells as previously described (17, 28). Analyses of *HTT* CAG repeat size in both *Htt*^Q111^ mice and in patient fibroblasts was performed by PCR using human-specific *HTT* primers and *Taq* PCR Core Kit with Q solution (Qiagen), as previously described (17, 43). The resulting FAM-labelled PCR products, encompassing the *HTT* CAG repeat, were resolved on the ABI3730xl automated DNA analyzer (Applied Biosystems), with either GeneScan 500-LIZ or 1200-LIZ internal size standards, and analyzed with GeneMapper v5 (Applied Biosystems). When necessary, to increase the PCR signal strength, up to three PCR replicates were combined and concentrated with MinElute PCR Purification Kit (Qiagen). CAG expansion indices were quantified in *Htt*^Q111^ mice from GeneMapper peak height data, taking into account only expansion peaks (*i.e*. larger than main allele) and using a 5% peak height threshold, as described previously (44). In the HD patient fibroblasts, CAG expansion was reflected in a shift of the entire CAG length distribution to greater repeat lengths over time, without any discernible impact on the shape of the distribution. To quantify repeat expansion, we determined the average gain of CAG units by calculating the difference in average CAG length before (*t* = 0, *n* = 4) and after the 6-week culture period (*t* = 6 weeks, *n* = 10). To determine average CAG length in an individual sample, we first applied a 20% peak height threshold to the distribution (i.e. only peaks with height ≥ 20% of tallest peak were used), then multiplied the frequency of each peak (peak height divided by the sum of all peak heights) by the corresponding CAG size, and finally summed these values.

### Treatment of *Htt*^Q111^ mice with *Mlh3* SSOs

Mouse-specific SSOs were designed to target the pre-mRNA of *Mlh3* exon 6 (Chr12:85,250,302-85,250,373; GRCm38/mm10): mMLH3ac5 (acceptor site) 5’-TCCACCTACAAAATAATCCAGGATT-3’ and mMLH3dr7 (donor site) 5’-AACTACAGACAGATACTTACCAGTA-3’. Oligonucleotides were obtained as “Vivo-Morpholinos” from Gene Tools, LLC (45). *Htt*^Q111^ mice (CAG 104-109; two males and two females per group) were treated with either 25 mg/kg SSOs (mMLH3ac5 and mMLH3dr7; 1:1 ratio), or PBS, by single tail vein injection of 200 μl. Treatment was initiated at 4 weeks of age and injections were performed three times per week (every 48-72hr). Weight measurements were taken at the beginning of each week to determine weekly weight-adjusted doses. Treatment was stopped at 12 weeks of age, with animal euthanasia and immediate dissection performed 24 hours after the final injection. Tissues were rapidly frozen using liquid nitrogen to preserve RNA integrity.

### *MLH3* splicing analysis

Frozen mouse tissues were mechanically homogenized prior to RNA extraction. Total RNA was extracted from *Htt*^Q111^ mouse tissues and HD patient cultured primary fibroblasts with TRIzol (Thermo Fisher Scientific) following the manufacturer’s protocol. Total RNA (250 ng) was used to generate cDNA with the High Capacity cDNA Reverse Transcription Kit following the manufacturer’s protocol (Thermo Fisher Scientific). Human *MLH3* splice variants were characterized by Reverse Transcription PCR (RT-PCR) as previously described (28). Mouse *Mlh3* splice variants were characterized by RT-PCR with primers mMLH3×3-3328F (5’-CCAGTGTTTGCTCGATACCC-3’) and mMLH3×11-4028R (5’-GGACAAGGACAGAGCTTCGA-3), at 59°C annealing temperature, for 33 cycles. PCR products were resolved by electrophoresis on a 1.4% agarose gel containing 1 μg/ml ethidium bromide. Real-time RT-PCR (RT-qPCR) characterization of *Mlh3* splicing in mouse tissues was performed with TaqMan Gene Expression Assays (Applied Biosystems) following the manufacturer’s protocol on the CFX96 Real-Time PCR Detection System (Bio-Rad). The assay used for the *Mlh3* splice variant 1 detects the splice junction between exon 6 and exon 7 (Mm00520819_m1, FAM-labeled probe). The assay for *Mlh3* splice variant 2, binds at the splice junction between exon 5 and exon 7 (custom design, FAM-labeled probe) and only binds when exon 6 is spliced out. Relative *Mlh3* splice variant levels were normalized to the endogenous mouse β-actin (*Actb*) reference gene (Mm00607939_s1, primer-limited, VIC-labeled probe) in duplex reactions.

### Treatment of HD patient primary fibroblasts with *MLH3* SSOs

Primary fibroblasts from a juvenile HD patient (GM09197; CAG ~180/18) (46) were obtained from the Coriell Institute for Medical Research. Cells were maintained in Dulbecco’s modified Eagle’s medium (DMEM) high glucose (Thermo Fisher Scientific) with 10% fetal bovine serum (FBS; Sigma) at 37°C in an atmosphere containing 5% CO_2_. To determine the ability of the SSOs to induce *MLH3* splice redirection in HD primary fibroblasts, confluent cell cultures of unaltered primary GM09197 cells were treated with a single dose of either 500 nM of *MLH3* SSOs (MLH3acr6 and MLH3dnr8, 1:1 ratio, 250 nM each) (28) or 500 nM of scrambled control oligonucleotide (5’-CCTCTTACCTCAGTTACAATTTATA-3’), alongside cells that were left untreated. Oligonucleotides were obtained as “Vivo-Morpholinos” from Gene Tools, LLC (45). Cells were treated for six hours in reduced-serum Opti-MEM media (Thermo Fisher Scientific). The media was then replaced with 20% FBS high glucose DMEM and cells were harvested for RNA isolation after 48 hours to analyze changes to *MLH3* mRNA splice variants. To test the long-term effect of *MLH3* splice redirection on *HTT* CAG expansion, GM09197 primary HD fibroblasts were first transduced with MSH3- and EGFP-expressing lentiviral particles (PNL-MSH3-IRES2EGFP construct), as previously described (27, 28), to induce CAG expansion. Cells were then grown to confluent cultures and treated twice weekly with 500 nM of *MLH3* SSOs, as descried above, or left untreated, for a total of six weeks. Cells were then harvested for DNA for *HTT* CAG instability analysis. A total of four wells were harvested immediately prior to treatment (*t* = 0) and a total of ten independent wells were harvested at the end of treatment (*t* = 6 weeks).

### Statistical analysis

Statistical analyses were performed using GraphPad Prism v.8. Comparison of *HTT* somatic CAG expansion indices between *Htt*^Q111^ mice with different *Mlh3*^DN^ genotypes and across different tissues was performed using two-way ANOVA, with Tukey multiple comparisons *post hoc* correction. Comparison of *HTT* somatic CAG expansion indices in *Htt*^Q111^ mice treated with *Mlh3* SSOs was performed using two-tailed unpaired *t* tests comparing *Mlh3* SSO-treated and PBS-treated mice for each tissue. Comparison of average *HTT* CAG gain in HD patient fibroblasts treated with *MLH3* SSOs was performed using one-way ANOVA (“*t* = 0” vs “*MLH3* SSOs, *t* = 6wks” vs “untreated, *t* = 6wks”), with Tukey multiple comparisons *post hoc* correction. Not significant, *p* > 0.05; *, *p* ≤ 0.05; **, *p* ≤ 0.01; ***, *p* ≤ 0.001; ****, *p* ≤ 0.0001.

## Results

### Mutation in the MLH3 endonuclease domain prevents HD CAG expansion in *Htt*^Q111^ mice

We previously demonstrated that MLH3 is required for somatic CAG repeat expansion in *Htt*^Q111^ knock-in mice (17). To test the hypothesis that MLH3 endonuclease activity is involved in repeat expansion, we used an *Mlh3* mutant mouse line (*Mlh3*^D1185N^, abbreviated to *Mlh3*^DN^) harboring a point mutation in the endonuclease domain (42). In these mice, <u>G</u>AC is changed to <u>A</u>AC in the genomic sequence to replace the aspartic acid “D” at amino acid 1185 with an asparagine “N” within the conserved <u>D</u>QHA(X)_2_E(X)_4_E motif (DQHAAHERIRLE in mouse MLH3) **(Figure 1A**). An analogous mutation in *S. cerevisiae* or human MLH3 eliminates its endonuclease activity (47–50). Importantly, this D to N substitution in purified mouse, human, and *S. cerevisiae* proteins did not alter the stability of MLH3 or its interaction with MLH1 (42, 50, 51). In addition, in the mouse there was no impact on the number or localization of MLH3 or MLH1 foci in spermatocytes or on the recruitment of factors required to initiate crossovers in meiosis (42). Molecular modeling of mouse MLH3 was consistent with the lack of impact of the D1185N substitution on the structure or stability of MLH3 (30). Therefore, the *Mlh3*^DN^ mutation in the mouse is predicted to disrupt MLH3 endonuclease function in the absence of significant effects on protein expression, stability, or interactions.

**Figure 1.**
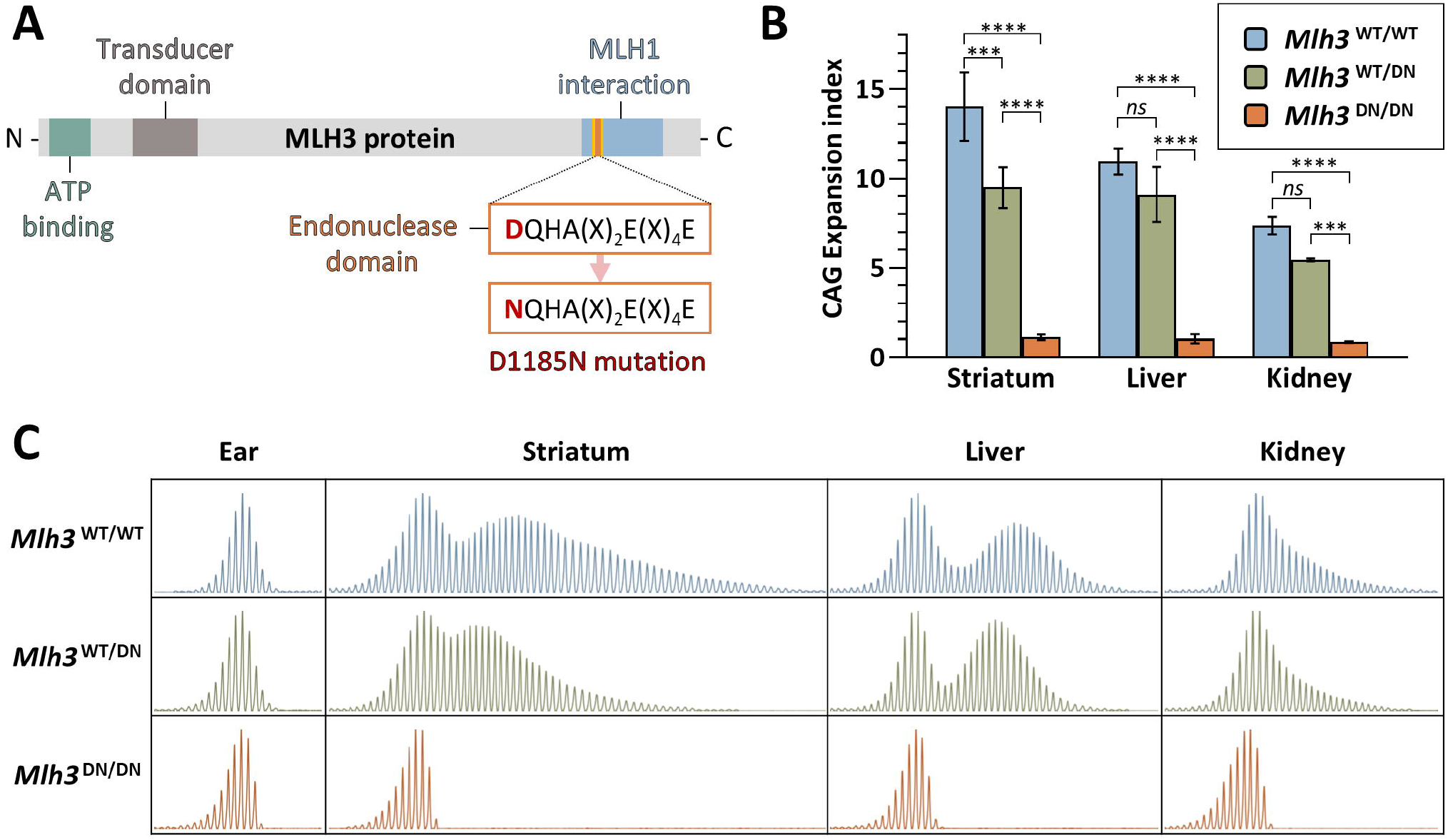
*MLH3* endonuclease domain mutation eliminates *HTT* CAG expansion in *Htt*^Q111^ mice. (**A**) Schematic representation of MLH3 protein and its functional domains, as well as mutation of the endonucleolytic motif in *Mlh3*^DN^ mice (adapted from Toledo *et al*. 2019 (42)). (**B**) Quantification of *HTT* CAG expansions in striatum, liver, and kidney at 6 months of age in *Htt*^Q111^ knock-in mice that are either wild type (*Mlh3*^WT/WT^, CAG 114-121), heterozygous (*Mlh3*^WT/DN^, CAG 116-121), or homozygous (*Mlh3*^DN/DN^, CAG 116-121) for the D1185N mutation shows that an intact MLH3 endonuclease domain is required for somatic CAG expansion. Striatum and liver, *n* = 3; kidney, *n* = 2-3; error bars represent standard deviation of the mean; two-way ANOVA (with genotype and tissue as variables) with Tukey multiple comparisons *post hoc* correction. Genotype effect: *ns*, not significant; ***, *p* ≤ 0.001; ****, *p* ≤ 0.0001. (**C**) Representative GeneMapper traces across different tissues at 6 months of age in *Htt*^Q111^ knock-in mice (CAG 121) with different *Mlh3* genotypes.

To determine the effect of *Mlh3*^DN^ allele on somatic *HTT* CAG expansion, we crossed *Mlh3*^DN^ mice (42) with *Htt*^Q111^ HD knock-in mice (41) to generate heterozygous *Htt*^Q111^ mice that were either wild type (*Mlh3*^WT/WT^), heterozygous (*Mlh3*^WT/DN^), or homozygous (*Mlh3*^DN/DN^) for the *Mlh3* D1185N point mutation. Analysis of CAG expansions in striatum, liver, and kidney – tissues previously shown to exhibit high levels of instability in *Htt*^Q111^ mice (9, 17) – revealed the absence of somatic expansions in *Mlh3*^DN/DN^ mice, both at three (**Figure S1**) and six months of age **(Figure 1B, C)**. Notably, the effect of the homozygous *Mlh3*^DN/DN^ mutation on CAG expansion was indistinguishable from that of a homozygous *Mlh3* null mutation (17). Quantification of repeat expansions (44) showed statistically significant reductions in the CAG expansion index in all tissues analyzed (striatum, liver, and kidney) of *Mlh3*^DN/DN^ mice compared to *Mlh3*^WT/WT^ mice **(Figure 1B** and **Figure S1**). There was a slight reduction in expansion index in the *Mlh3*^WT/DN^ mice, consistent at both ages and in all tissues analyzed, though only reaching statistical significance in 6-month striata **(Figure 1B** and **Figure S1**). The dramatic effect of this mutation, phenocopying the *Mlh3* null (17), indicates that an intact endonuclease domain is required for somatic *HTT* CAG repeat expansion.

### *In vivo Mlh3* splice redirection suppresses CAG expansion in *Htt*^Q111^ mice

Having demonstrated that the MLH3 endonuclease domain was critical for somatic *HTT* CAG expansion in *Htt*^Q111^ mice, we were interested in testing the potential of an *Mlh3* splice redirection approach, previously used in FRDA cell-based models and patient cells (28), to slow CAG expansion in the same HD mice. Mice express predominantly one endogenous *Mlh3* mRNA splice variant in which exon 6, the homologous exon to human *MLH3* exon 7, encodes the endonuclease domain **(Figure 2A** and **Figure S2C**). For this study, mouse-specific vivo-morpholino-based SSOs were designed to bind both the acceptor and donor splice sites of *Mlh3* exon 6 **(Figure 2A** and **Figure S2B**), as previously reported for human *MLH3* (28). Starting at 4 weeks of age, heterozygous *Htt*^Q111^ mice (*n* = 4) were treated with 25 mg/kg of *Mlh3* SSOs (or PBS) by tail vein injection every 48-72 hours, until they reached 12 weeks of age **(Figure 2B**). *Htt*^Q111^ tissues were harvested 24 hours after the final injection and *Mlh3* splice variants were characterized by RT-PCR analysis in liver and kidney **(Figure 2C)**, two peripheral tissues in which CAG expansion was shown to be dependent on an intact MLH3 endonuclease domain **(Figure 1B, C**) and which can be effectively targeted with systemic delivery of vivo-morpholinos (45). We analyzed *Mlh3* splice variants using RT-PCR, generating a 720 bp product for the exon 6-containing splice variant 1 (Var1) and a 648 bp product for *Mlh3* splice variant 2 (Var2), which lacks exon 6 (**Figure S2C**) and is the mouse analog of human *MLH3* splice variant 2. As expected, all four PBS-treated mice showed Var1 only in both kidney and liver, while all four *Mlh3* SSO-treated mice showed successful splice redirection favoring Var2 in the kidney, with much lower, yet still detectable, Var2 levels in the livers of two treated mice **(Figure 2C**). We also established a TaqMan-based real-time RT-PCR assay for quantitative analysis of *Mlh3* splice variants and increased sensitivity (**Figure S3**). Using this more sensitive assay, only Var1 was detected in PBS-treated mice, whereas variable expression of Var2 was detected in all SSO-treated samples, with the global pattern of Var1 and Var2 expression in the two tissues consistent with levels detected in the gel-based RT-PCR assay (**Figure S3**). The higher levels of Var2 in the kidney relative to the liver may reflect better cellular SSO uptake in the kidney than in the liver, or may be the result of SSOs being actively removed from circulation by the kidney for excretion, limiting SSO availability to other peripheral tissues. In summary, we found that *Mlh3* SSO-mediated splice redirection to exclude the exon encoding the endonuclease domain of MLH3 can be successfully achieved *in vivo* in a mouse model of HD.

**Figure 2.**
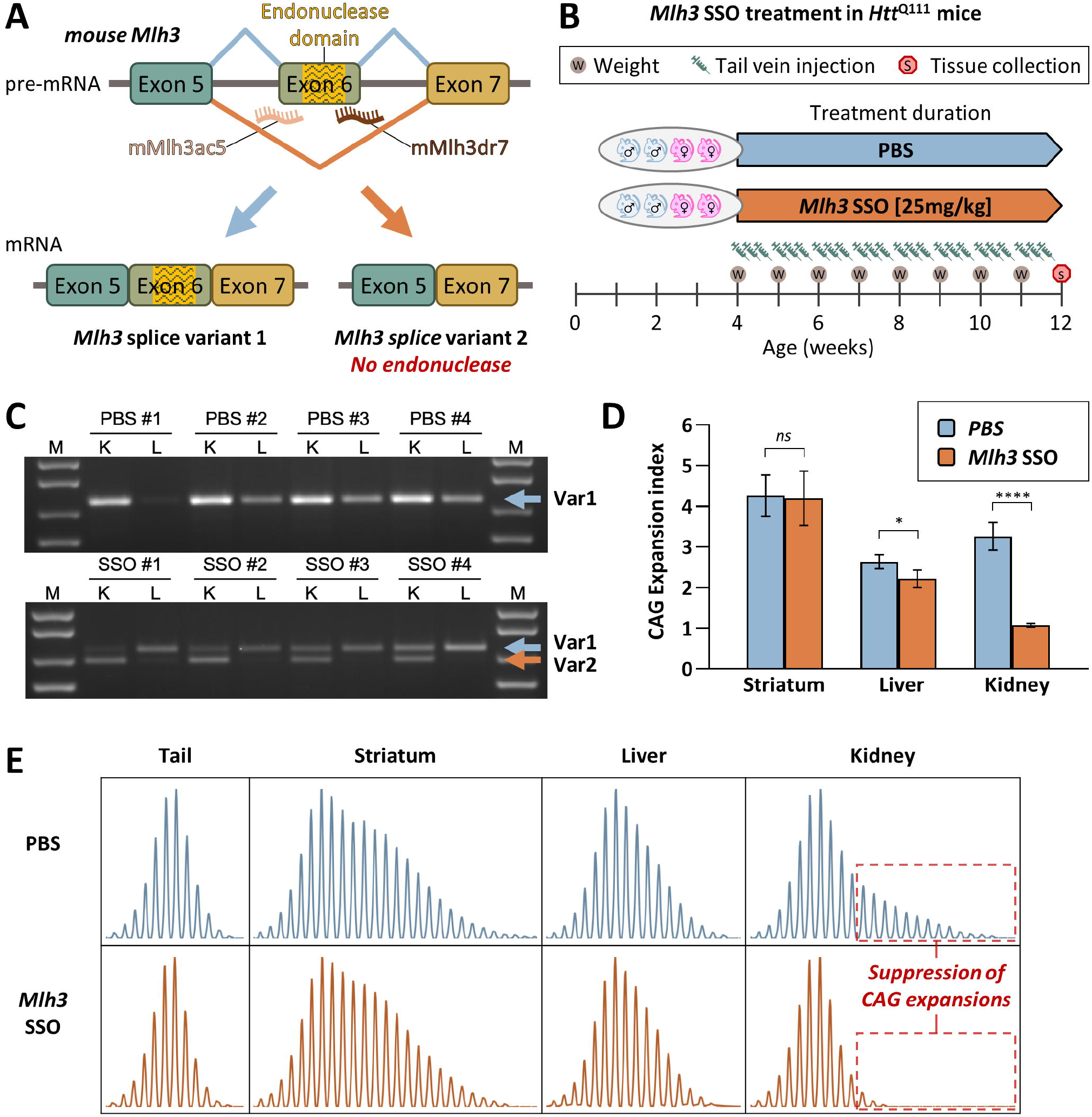
Successful *in vivo Mlh3* splice redirection and suppression of somatic *HTT* CAG expansion in *Htt*^Q111^ mice by systemic *Mlh3* SSO treatment. (**A**) Schematic representation of mouse *Mlh3* splice redirection with splice switching oligonucleotides (SSOs). SSOs were designed to bind the intron–exon junctions in mouse *Mlh3* pre-mRNA at either the splice acceptor (mMlh3ac5) or donor (mMlh3dr7) regions of the endonuclease-coding exon 6, which is analogous to the human MLH3 exon 7, inducing exon skipping and preferential production of *Mlh3* splice variant 2, which lacks the endonuclease domain. (**B**) Summary of *Mlh3* SSOs systemic treatment in HD mice. *Htt*^Q111^ mice (two males and two females per group) were treated with either 25 mg/kg *Mlh3* SSOs (mMLH3ac5 and mMLH3dr7, 1:1 ratio; CAG 106-109), or PBS (CAG 104-108), by tail vein injection. Treatment was initiated at 4 weeks of age and injections were performed three times per week (every 48-72hr). Weight measurements were taken at the beginning of each week to determine weekly weight-adjusted doses. Treatment was stopped at 12 weeks of age, with tissue collection performed 24 hours after the final injection. (**C**) RT-PCR analyses of *Mlh3* mRNA splice variants in 12-week-old *Htt*^Q111^ mice, following treatment with *Mlh3* SSOs, reveals successful splice redirection from splice variant 1 (Var1) to variant 2 (Var2) in the kidney (K), and to a lesser extent in the liver (L). (**D**) Quantification of CAG expansion indices reveals statistically significant reductions in somatic *HTT* CAG expansion in kidney and, to a lesser extent, in liver of mice that received *Mlh3* SSO treatment when compared to PBS. *n* = 4; error bars represent standard deviation of the mean; 2-tailed unpaired *t* test: *ns*, not significant; *, *p* ≤ 0.05; ****, *p* ≤ 0.0001. (**E**) Representative GeneMapper traces of somatic *HTT* CAG repeat size distributions across different tissues at 12 weeks of age in *Htt*^Q111^ knock-in mice (CAG 108), following PBS or *Mlh3* SSO treatment.

To test whether such *Mlh3* splice redirection impacted somatic CAG expansion, we performed qualitative and quantitative analyses of CAG repeat length profiles from the same *Htt*^Q111^ mice (*n* = 4) treated with either *Mlh3* SSOs or PBS **(Figure 2D, E**). When compared to the PBS-treated mice, the *Mlh3* SSO-treated mice displayed significantly less somatic CAG expansion in the kidney (*p* ≤ 0.0001), and to a lower extent in the liver (*p* ≤ 0.05). These effects are consistent with the different degrees of splice redirection achieved in the two tissues. As expected, *Mlh3* SSO treatment had no effect on CAG expansion in the striatum, consistent with the inability of these SSOs to efficiently cross the blood brain barrier (45). As *Mlh3* splice redirection from Var1 to Var2 increases the proportion of mRNAs encoding an endonuclease-lacking MLH3 protein, these data support our prior results in *Mlh3*^DN^ mice showing that an intact endonuclease domain is required for somatic *HTT* CAG expansion. Further, these results demonstrate that a pharmacological splice redirection strategy can be used *in vivo* to impact a function of MLH3 that is critical for somatic CAG expansion.

### *MLH3* splice redirection suppresses CAG expansion in HD patient-derived fibroblasts

Alternative splicing of human *MLH3* pre-mRNA results in endogenous expression of two predominant splice variants: variant 1 (Var1), containing the exon7-encoded endonuclease domain, and variant 2 (Var2), lacking this domain **(Figure 3A**, **Figure S2C** and **Figure S4**). We previously showed that *MLH3* splice redirection from Var1 to Var2 can be achieved in FRDA patient primary fibroblasts using a combination of SSOs targeting the exon 7 splice acceptor (MLH3acr6) and donor (MLH3dnr8) sites (28). Prolonged treatment with these *MLH3* SSOs resulted in suppression of GAA repeat expansion in FRDA cell-based models and patient cells (28). With the goal of testing this strategy in HD, we used primary fibroblasts derived from a juvenile HD patient (GM09197; CAG ~180/18) (46) with a long repeat tract, predicted to allow repeat expansion in cell culture within a reasonable time frame (27, 52). We first tested our ability to induce *MLH3* splice redirection following 48 hour treatment with 500 nM of *MLH3* SSOs (MLH3acr6 and MLH3dnr8, 1:1 ratio, 250nM each) compared to treatment with 500 nM of a control vivo-morpholino oligonucleotide with a scrambled sequence or no treatment **(Figure 3B**). RT-PCR analysis of the *MLH3* splice variants generates a 434 bp product for the full-length Var1 and a 362 bp product for Var2. RT-PCR analysis detects *MLH3* Var1 as the primary isoform expressed in GM09197 fibroblasts, while treatment with *MLH3* SSOs resulted in full redirection of splicing towards Var2 **(Figure 3B**). No impact on *MLH3* splicing was detected following treatment with scrambled control oligonucleotides, where splice variants were identical to untreated cells. We then determined whether splice redirection of *MLH3* was capable of slowing CAG expansion in these cells over time. In our studies on FRDA primary fibroblast cells, we found that the GAA repeat tract was stable due to low levels of expression of MSH3, one of the key factors that contribute to both GAA and CAG repeat expansion (27, 53). Therefore, we induced CAG expansion in the HD fibroblasts via ectopic expression of MSH3 using lentiviral vector delivery as previously described for FRDA fibroblasts (27, 28). These MSH3-expressing fibroblasts were then grown to confluent cultures to prevent replication, before initiating SSO treatment. Prior to SSO treatment, genomic DNA was isolated from untreated cells (*t* = 0; *n* = 4) to determine the starting average CAG repeat size. Cells were then treated with 500 nM *MLH3* SSOs twice-weekly, or left untreated, for a total of six weeks (*t* = 6wks) **(Figure 3C**). Quantitative GeneMapper analysis revealed an overt shift of the entire population of CAG alleles towards larger lengths in untreated HD fibroblasts (*t* = 6wks), relative to the starting population (*t* = 0), reflected by an increase in average CAG lengths of 3.0 repeats (*p* ≤ 0.0001) **(Figure 3D, E** and **Figure S5**). Treatment with *MLH3* SSOs for 6 weeks significantly suppressed CAG expansion, with cells showing only a modest and non-statistically significant increase in average CAG length of 0.6 repeats during this period (*p* ≤ 0.0001, when compared with untreated cells, *t* = 6 weeks; *p* = 0.3120 when compared with *t* = 0) **(Figure 3D, E** and **Figure S5**). These data further support our prior results in *Mlh3*^DN^ and *Mlh3* SSO-treated *Htt*^Q111^ mice, suggesting that the MLH3 endonuclease domain is essential for somatic *HTT* CAG expansion in HD patient cells, and that the use of *MLH3* SSOs constitutes a viable and effective strategy to pharmacologically target this pathway in HD patients.

**Figure 3.**
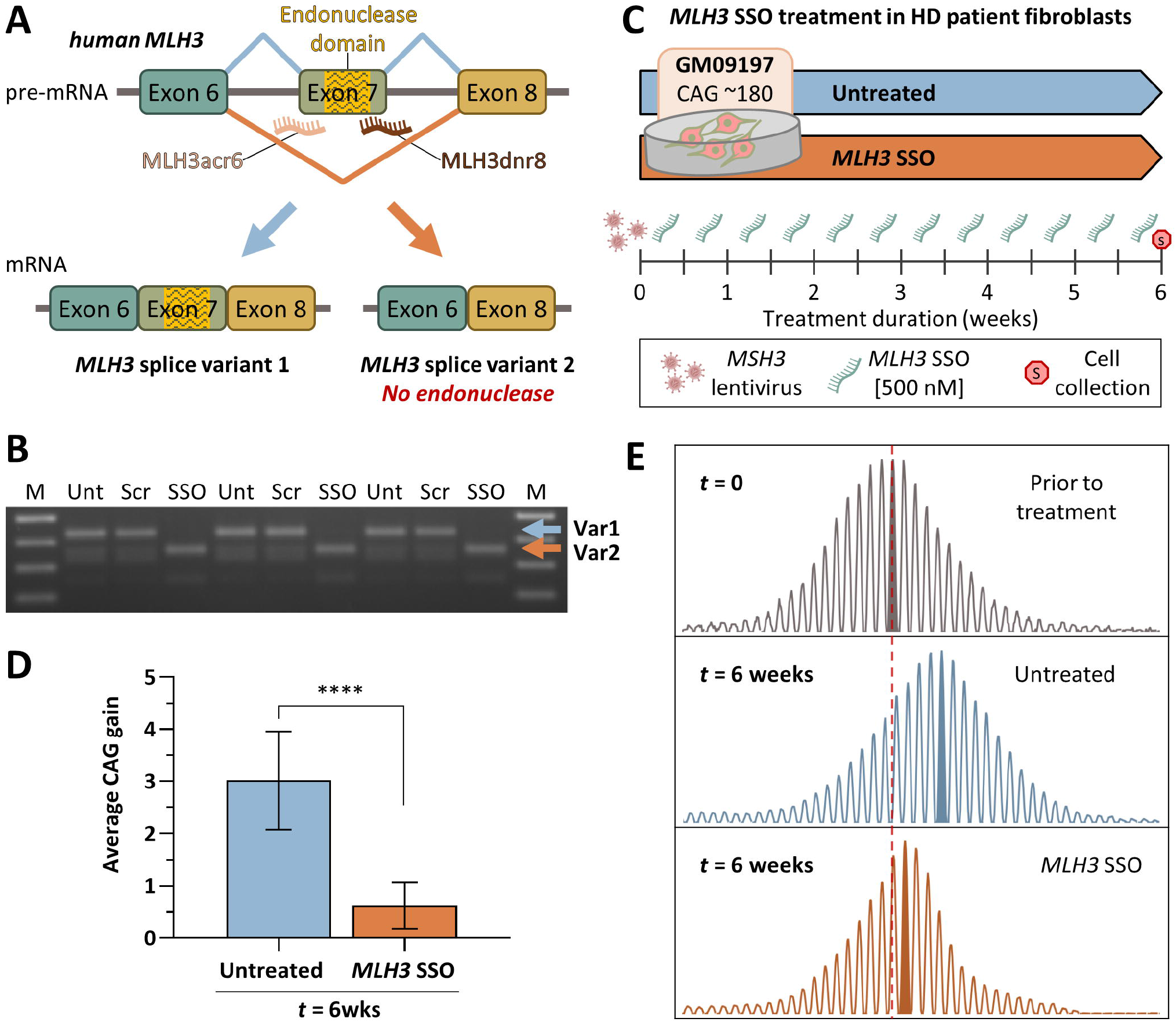
Successful *MLH3* splice redirection and suppression of *HTT* CAG expansion in HD patient-derived fibroblasts following treatment with *MLH3* SSOs. (**A**) Schematic representation of human *MLH3* splice redirection with splice switching oligonucleotides (SSOs) (adapted from Halabi *et al*. 2018 (28)). SSOs were designed to bind the intron–exon junctions in human *MLH3* pre-mRNA at either the splice acceptor (MLH3ac6) or donor (MLH3dnr8) regions of the endonuclease-coding exon 7, inducing exon skipping and preferential production of *MLH3* splice variant 2, which lacks the endonuclease domain. (**B**) RT-PCR analyses of *MLH3* mRNA splice variants in HD patient-derived primary fibroblasts (GM09197, CAG ~180/18) following treatment with 500 nM of either scrambled control vivo-morpholino oligonucleotide (Scr) or *MLH3* SSOs (SSO) for 48 hours, alongside untreated controls (Unt). Splice redirection from predominantly splice variant 1 (Var1) to predominantly splice variant 2 (Var2), which lacks exon 7 and the endonuclease domain, was efficiently achieved with *MLH3* SSOs treatment. No splice redirection was observed in scrambled oligo-treated cells, which were identical to untreated cells. (**C**) In order to induce CAG expansion in HD patient fibroblasts (GM09197, CAG ~180/18), cells were initially treated with lentiviral particles for stable ectopic expression of MSH3. Cells were then cultured to confluency, to inhibit replication, and treated twice weekly with 500 nM of *MLH3* SSOs (MLH3acr6 and MLH3dnr8, 1:1 ratio, 250 nM each), or left untreated, for a total of six weeks when cells were harvested for *HTT* CAG instability analysis. (**D**) Quantification of average *HTT* CAG gain during the 6 week-long treatment (relative to average CAG prior to treatment, “*t* = 0”, *n* = 4) reveals a potent inhibition of CAG expansion by *MLH3* SSOs (average gain of 0.6 CAGs, *n* = 10), when compared to untreated cells (average gain of 3.0 CAGs, *n* = 10). Error bars represent standard deviation of the mean; one-way ANOVA with Tukey multiple comparisons *post hoc* correction; ****, *p* ≤ 0.0001. (**E**) Representative GeneMapper traces of *HTT* CAG repeat size distributions from GM09197 HD patient fibroblasts before (*t* = 0) and after (*t* = 6 weeks) treatment with *MLH3* SSOs (CAG gain = 0.7), or no treatment (CAG gain = 3.6).

## Discussion

Somatic expansion of the *HTT* CAG repeat is thought to drive the rate of HD pathogenesis. Repeat expansion is dependent on proteins in the MMR pathway and, therefore, these present potential therapeutic targets. Insight into specific functions or activities of MMR proteins that modify repeat expansion will provide possible opportunities for therapeutic intervention. Here, we provide evidence, using two distinct, yet complementary approaches, that the endonuclease domain of MLH3 is required for somatic *HTT* CAG expansion. First, a point mutation in a conserved endonuclease domain motif abolishes CAG expansion in *Htt*^Q111^ mouse tissues, having an effect indistinguishable from a *Mlh3* null mutation. Second, CAG expansion both in *Htt*^Q111^ mice and in HD patient-derived cells is reduced by modulating *Mlh3*/*MLH3* splicing in a manner that excludes the exon encoding the endonuclease motif.

The endonuclease domain of MLH3 is encompassed in the C-terminal domain of the protein, which interacts with MLH1 (54). As described above, whereas the D>N substitution in the first amino acid of the DQHA(X)2E(X4)E motif eliminates MLH3 endonuclease activity (47–50), there is no evidence that this mutation alters the stability of MLH3 or its interaction with MLH1 (30, 42, 50, 51). We have been unable to directly determine MLH3 protein levels in our study due to the lack of availability of suitable antibodies to detect MLH3, which is present at very low levels in most tissues (28, 38). More specifically, we could not detect MLH3 in mouse tissues using the same antibody used to detect MLH3 in mouse spermatocytes of *Mlh3*^DN^ mice, where this protein is relatively highly expressed (42). Importantly, we find that heterozygous or homozygous *Mlh3*^DN^ alleles have impacts on CAG expansion comparable to heterozygous or homozygous *Mlh3* null alleles, respectively (17). This indicates that the *Mlh3*^DN^ mutation would need to dramatically reduce MLH3 protein levels in order to have an impact on CAG expansion that phenocopies the null allele. Given existing data in the literature (30, 42, 50, 51), this scenario appears highly improbable. Further, the comparable effects of the *Mlh3*^DN^ and *Mlh3* null alleles argue against a dominant negative effect of the *Mlh3*^DN^ mutation. It is therefore likely that the absence of *HTT* CAG expansion in *Htt*^Q111^ mice harboring the *Mlh3* D1185N mutation is due to loss of MLH3-dependent endonuclease activity.

The impact of *Mlh3/MLH3* splice redirection that excludes the DQHA(X)2E(X4)E motif on *HTT* CAG expansion is also consistent with a requirement for MLH3-dependent endonuclease activity, although we cannot rule out the possibility that deletion of the endonuclease domain-encoding exon in MLH3 due to splice redirection has additional effects on MLH3 stability or function. Published data in this regard provide evidence that the human MLH3Δ7 isoform can interact with MLH1 in yeast 2-hybrid (54) and GST pull down experiments (55), and is recruited to repair foci in response to DNA damage in the same way as the full length MLH3 (56), implying normal protein interactions. However, one study found that MLH3Δ7 was unable to interact with MLH1 in a mammalian 2-hybrid assay (57). Although the endonuclease domain is encompassed in MLH3 and in the homologous PMS2 protein, endonuclease activity is dependent on the interaction of these proteins with MLH1 (MLH1-MLH3 = MutLγ dimer; MLH1-PMS2 = MutLα dimer), and structural analysis has revealed that MLH1 contributes an amino acid residue to the endonuclease active site of *S. cerevisiae* MutLα (58, 59). Thus, destabilizing this interaction would also be predicted to abrogate endonuclease activity. Overall, our data from orthogonal genetic and pharmacological-based approaches together provide strong support for a critical role of the MLH3 endonuclease in driving somatic expansion of the *HTT* CAG repeat.

As described previously, exclusion of *MLH3* exon 7 slowed GAA repeat expansion in FRDA cell models and in FRDA patient cells (28), and more recently, the same *Mlh3* D1185N point mutation was found to eliminate expansion of the *FXN* CGG repeat in a FXD mouse embryonic stem cell model (30). Together, data indicate that the MLH3 endonuclease domain is critical for somatic expansion of different trinucleotide repeat sequences, with implications for cross-disease applicability of therapeutic approaches that target this activity. Recent biochemical studies in cell-free systems have suggested a plausible mechanism by which MLH3 endonuclease activity might lead to repeat expansion (50). These studies provided support that MutLγ cleaves a closed circular plasmid harboring a CAG- or CTG-containing loop on the loop-lacking strand, with repair leading to loop incorporation, effectively corresponding to an expansion event. This observation distinguished the MutLγ endonuclease from the MutLα endonuclease, where MutLα was able to cleave on both the loop-containing and loop-lacking strands, potentially leading to both deletion and expansion events (60). This specific strand-directed activity of MutLγ may, in part, explain the necessity for MLH3 in the somatic expansion of several disease-associated repeats (17, 28, 31). This is in contrast to PMS2, whose knockout can fully (32) or partially (61) suppress expansion, or can promote expansion (26, 28), depending on the repeat sequence and/or biological system. Understanding the role played by PMS2 endonuclease in somatic repeat instability *in vivo* would also be of significant interest.

The fact that *MLH3* splice variants that both include or exclude the endonuclease domaincontaining exon 7 occur naturally in humans indicates that the relative levels of these variants might have biological relevance. As exon 7 exclusion resulted in reduced CAG expansion, this prompted us to examine whether the levels of *MLH3* Var2 might correlate with tissue-specific expansion propensities of the *HTT* CAG repeat (43). Examination of *MLH3* Var1 and Var2 mRNA expression in the GTEx database (**Figure S4**) did not reveal any obvious correlation with levels of CAG expansion in tissues; where forebrain regions such as the cortex and basal ganglia tend to have high instability, cerebellum exhibits low instability, and in the periphery, liver exhibits high instability (43). While more refined cell type-based analyses would be needed to better understand relationships between CAG instability and *trans*-modifying factors, these initial observations do not indicate any obvious relationship between naturally occurring *MLH3* Var2 and tissue-specific CAG expansion.

How is the MutLγ endonuclease activated to promote repeat expansion? MutLγ endonuclease is also required in canonical MMR in *S. cerevisiae* and in meiosis (42, 51), however the mechanism(s) of endonuclease activation are not yet well understood. In reconstituted systems, MutLγ can bind directly to DNA oligonucleotides, including those containing loop or branched structures, but does not cleave these structures (47–49, 62). This, and the further observation that larger DNA molecules are better substrates for the MutLγ endonuclease suggested that MutLγ binds cooperatively to DNA and that higher order polymers are required for endonuclease activation (62). Interestingly, the nicking activity of a 50:50 mixture of wild-type Mlh1-Mlh3 and Mlh1-Mlh3D523N dimers (*S. cerevisiae* D523N is orthologous to the D1185N mouse mutation) was the same as that elicited by 100% wild-type Mlh1-Mlh3, consistent with the idea that a critical level of DNA binding is needed for endonuclease activity (62). It is unknown whether MutLγ polymers are required *in vivo*, but this observation suggests one possible explanation for the relatively minor impact on CAG instability in heterozygous *Mlh3*^WT/DN^ mice. In contrast, it appears that when the endonuclease-competent MLH3 is reduced below 50%, as in the kidney of *Htt*^Q111^ mice treated with *Mlh3* SSOs (**Figure S3**), the effect on expansion is more pronounced **(Figure 2C, E**).

In reconstituted systems, both human and *S. cerevisiae* MutLγ are stimulated by MutSβ (human MSH2-MSH3 or *S. cerevisiae* Msh2-Msh3 respectively), potentially via direct interaction of these two complexes (47, 50, 63), but human MutLγ was not stimulated by MSH2-MSH6 (MutSα) (63). As MSH3, but not MSH6, is required for trinucleotide repeat expansion (16), MutSβ-stimulation of MutLγ endonuclease may be integral to repeat expansion *in vivo*. Other protein-protein interactions have been implicated in MutLγ endonuclease activation (63–65) and, interestingly, *S. cerevisiae Mlh3* separation of function mutations provided evidence that distinct protein interactions may be required for MutLγ endonuclease activity in MMR and in meiosis (63, 64). Activity appears to be variably stimulated by proliferating cell nuclear antigen (PCNA) and replication factor C (RFC), depending on the assay system (47, 50, 63, 65). Studies in *S. cerevisiae* have also indicated that *Mlh3* endonuclease creates DNA breaks in the presence of RNA/DNA hybrids (R-loops) that can lead to increased instability, suggesting a link with transcription (66), though the relevance to repeat expansion *in vivo* is unclear. Future biochemical and genetic studies will be needed to dissect the mechanism of MutLγ endonuclease activation in its repeat expansion-promoting function.

Is *MLH3* a good therapeutic target for HD? In contrast to other MMR pathway genes (*MSH3, MLH1 PMS2, PMS1*), *MLH3* is not currently validated as a disease modifier in human GWAS (14). This may relate to the effect size of a putative modifier SNP(s) and the power to detect a significant effect on a clinical endpoint with the current patient sample size. Nevertheless, the following provide support for *MLH3* as a viable therapeutic target in HD: (1)*MLH3* is part of a pathway involved in disease modification that likely acts via somatic repeat expansion, (2) *MLH3* is essential for somatic CAG repeat expansion in HD mice (17), (3) Functional MLH3 promotes *HTT* CAG expansion in HD patient cells (this study), and (4) *MLH3* genetic variation is associated with somatic *HTT* CAG expansion measured in HD patient blood DNA (13). From a safety standpoint, consistent with a minor role of MLH3 in MMR, only a handful of studies have reported associations of *MLH3* mutations with cancer, however the pathogenicity of these mutations and their clinical significance are still unclear (67–70). Further, the genome Aggregation Database (gnomAD) reports an extremely low probability for intolerance to loss of function mutations for *MLH3* (pLI = 0.00) (71). Therefore, MLH3 appears to be a promising candidate as a safe and effective therapeutic target.

Here, we demonstrate applicability to HD of one possible therapeutic approach targeting the critical endonuclease domain of MLH3 using SSOs to redirect *MLH3* splicing. Future studies will be needed to demonstrate the efficacy of *MLH3* splice redirection in the brain. The splice redirection approach has the advantage of specifically altering the MLH3 endonuclease whilst leaving intact the PMS2 endonuclease that is critical to the major MMR activity in cells. Further insights into distinguishing features of these endonucleases may provide additional opportunities for specific inactivation of the MLH3 endonuclease. An alternative oligonucleotide-based approach of simply reducing *MLH3* expression would also be indicated based on genetic data in mice. In contrast to splice redirection, this strategy would reduce or eliminate all MLH3 function(s). Indeed, by some measures of MLH3 activity, the *Mlh3*^DN/DN^ is indistinguishable from *Mlh3*^WT/WT^, suggesting that the splice redirected MLH3 protein would also retain such nuclease-independent activities (42). Therefore, additional understanding of the cellular roles of the MLH3 endonuclease, and of the endonuclease-lacking variant, will help to inform on optimal therapeutic approaches applicable to both HD and other repeat expansion diseases.

## Acknowledgments

We would like to thank Tammy Gillis and Jacqi Siciliano of the MGH Mission Driven Service Core for DNA Fragment Analysis.

