## Supplemental Figures for "Somatic CAG expansion in Huntington’s disease is dependent on the MLH3 endonuclease domain, which can be excluded via *MLH3* splice redirection to suppress expansion"

**Figure S1. *MLH3* endonuclease domain mutation eliminates *HTT* CAG expansion in *Htt*$^{Q111}$ mice at 3 months of age.**

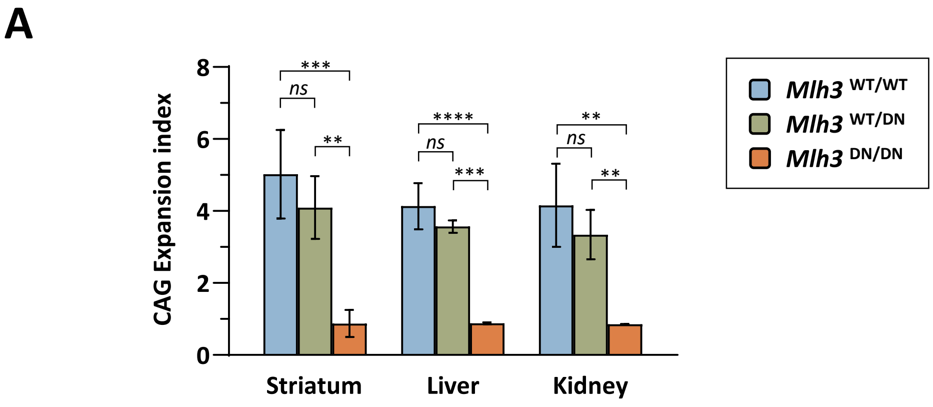


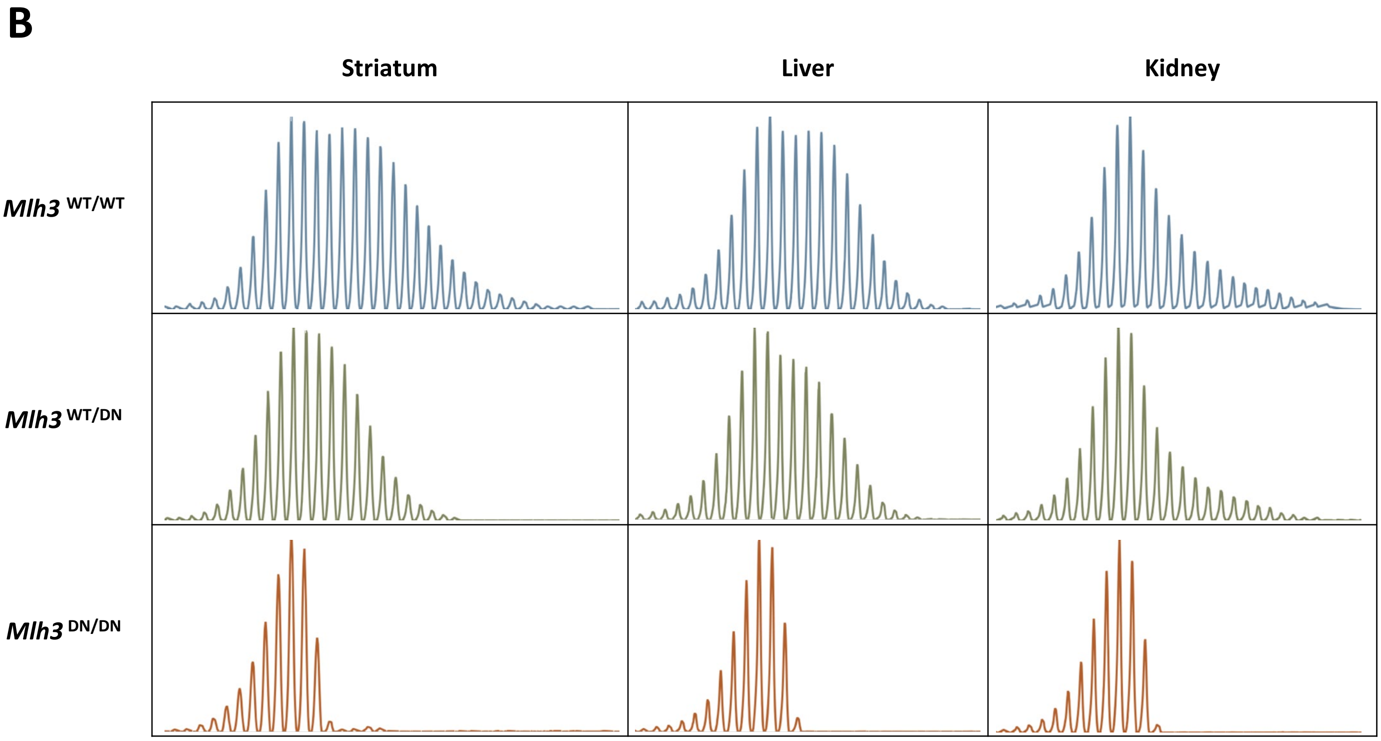


(**A**) Quantification of *HTT* CAG expansions in striatum, liver, and kidney at 3 months of age in *Htt*$^{Q111}$ knock-in mice that are either wild type (*Mlh3*$^{WT/WT}$, CAG 115-122), heterozygous (*Mlh3*$^{WT/DN}$, CAG 115-121), or homozygous (*Mlh3*$^{DN/DN}$, CAG 116-121) for the D1185N mutation. Striatum, *n* = 5-6; liver, *n* = 5; kidney, *n* = 4-5; error bars represent standard deviation of the mean; two-way ANOVA (with genotype and tissue as variables) with Tukey multiple comparisons *post hoc* correction. Genotype effect: *ns*, not significant; **, $p \leq 0.01$; ***, $p \leq 0.001$; ****, $p \leq 0.0001$. (**B**) Representative GeneMapper traces across different tissues at 3 months of age in HttQ111 knock-in mice with different Mlh3 genotypes. *Mlh3*$^{WT/WT}$, CAG 119, *Mlh3*$^{WT/DN}$ and *Mlh3*$^{DN/DN}$, CAG 118.

**Figure S2. Mouse *Mlh3* exon 6 splice exclusion results in a variant that lacks the endonuclease domain but otherwise maintains the same translational frame.**

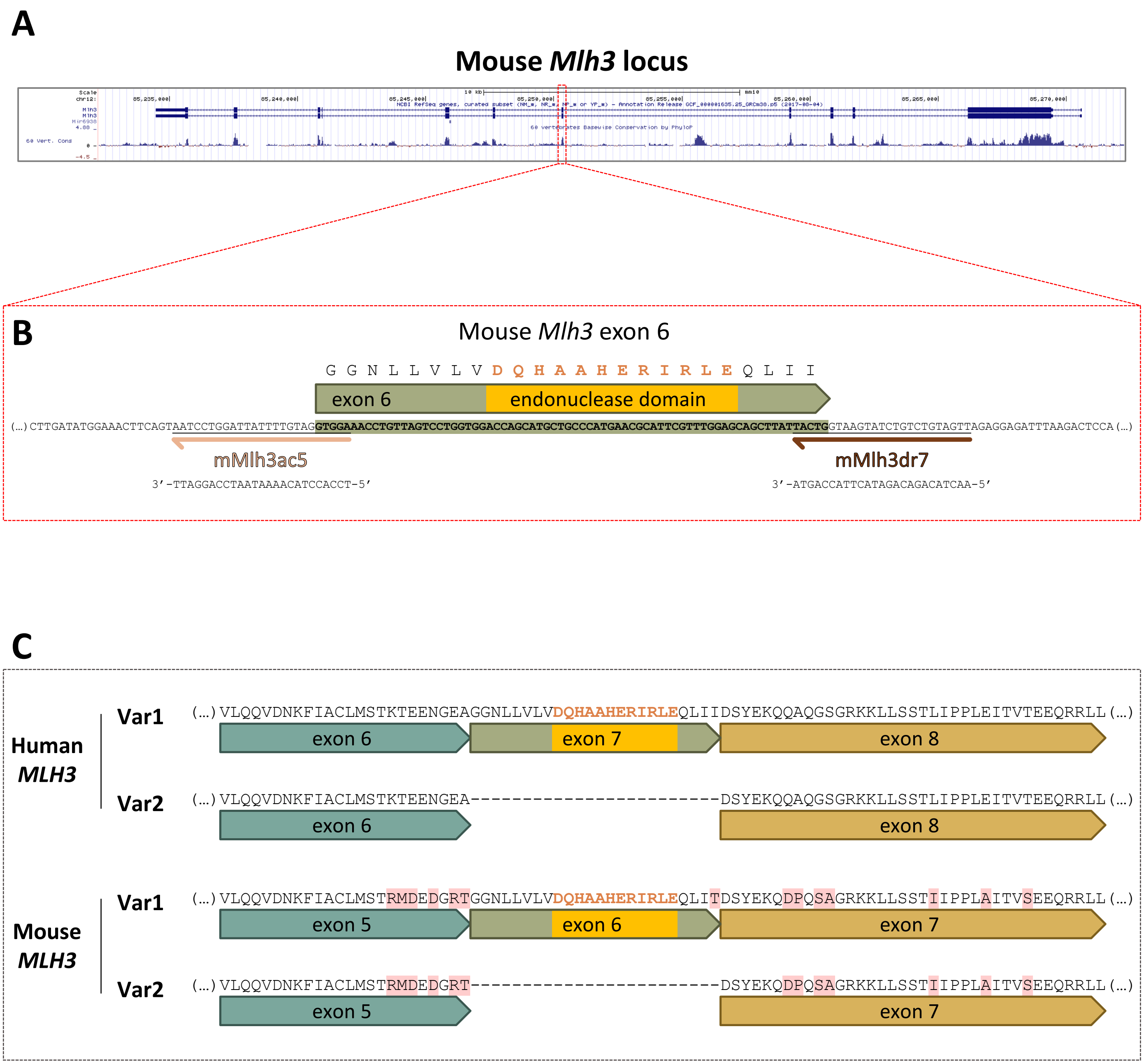


(**A**) Mouse *Mlh3* locus: chr12:85,232,251-85,272,250; GRCm38/mm10 assembly; NCBI RefSeq genes, curated subset; UCSC Genome Browser. (**B**) Splice redirection of mouse *Mlh3* using SSOs (mMlh3ac5 and mMlh3dr7) to exclude exon 6, which encodes the endonuclease domain (DQHAAHERIRLE). (**C**) Alignment of MLH3 protein isoforms encoded by human and mouse splice variants 1 and 2 (Var1 and Var2, respectively).

**Figure S3. Relative expression of mouse *Mlh3* splice variants 1 and 2 after splice redirection with SSOs.**

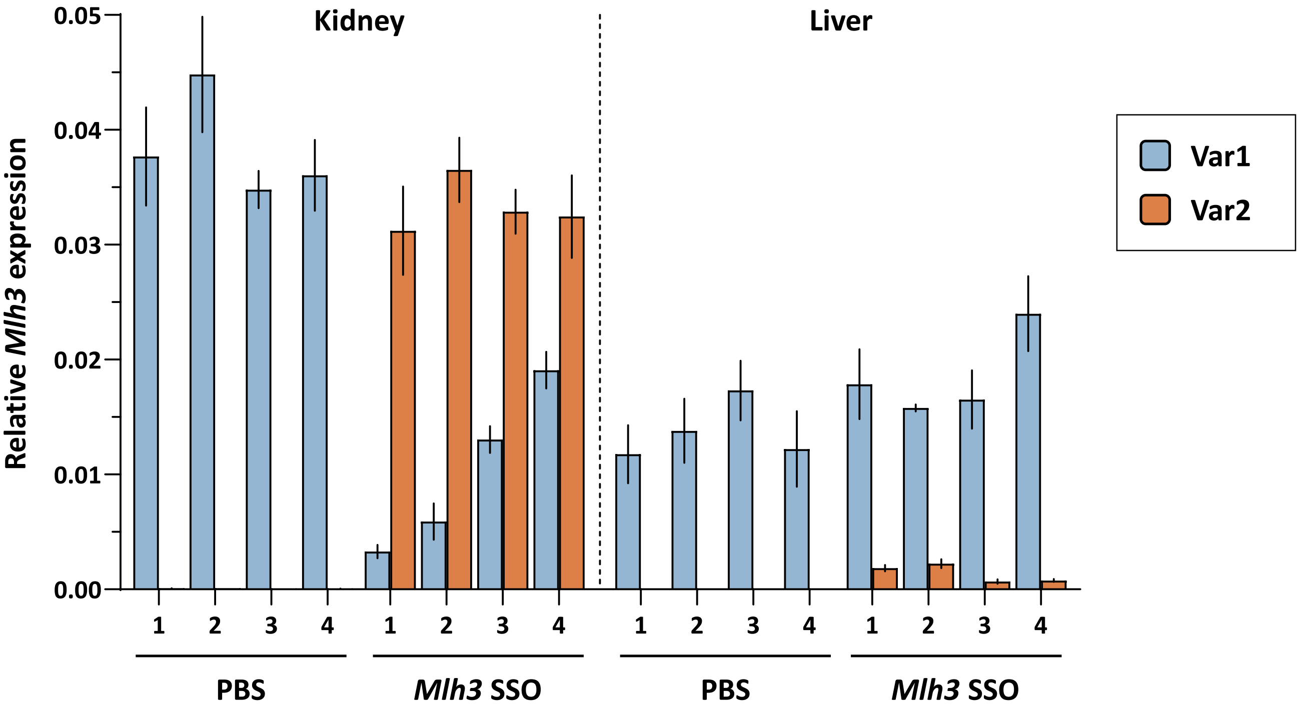


Quantification of *Mlh3* splice variants by RT-qPCR in *Htt*$^{Q111}$ mice (CAG 104-109) at 12 weeks of age, following 8-week-long treatment with either 25 mg/kg *Mlh3* SSOs (mMLH3ac5 and mMLH3dr7, 1:1 ratio) or PBS. Tissue collection was performed 24hrs after the final tail-vein injection. *Mlh3* SSO treatment successfully induced splice redirection from variant 1 (Var1) to variant 2 (Var2) in the kidney and, to a lesser degree, in the liver. *Mlh3* expression is relative to endogenous mouse β-actin (*Actb*). Since *Actb* mRNA levels can vary across different mouse tissues, relative *Mlh3* expression cannot be directly compared between kidney and liver.. Biological replicates, *n* = 4 (two males and two females per group); technical replicates, *n* = 3; error bars represent standard deviation of the mean.

**Figure S4. Expression of human *MLH3* splice variants 1 and 2 across multiple tissues.**

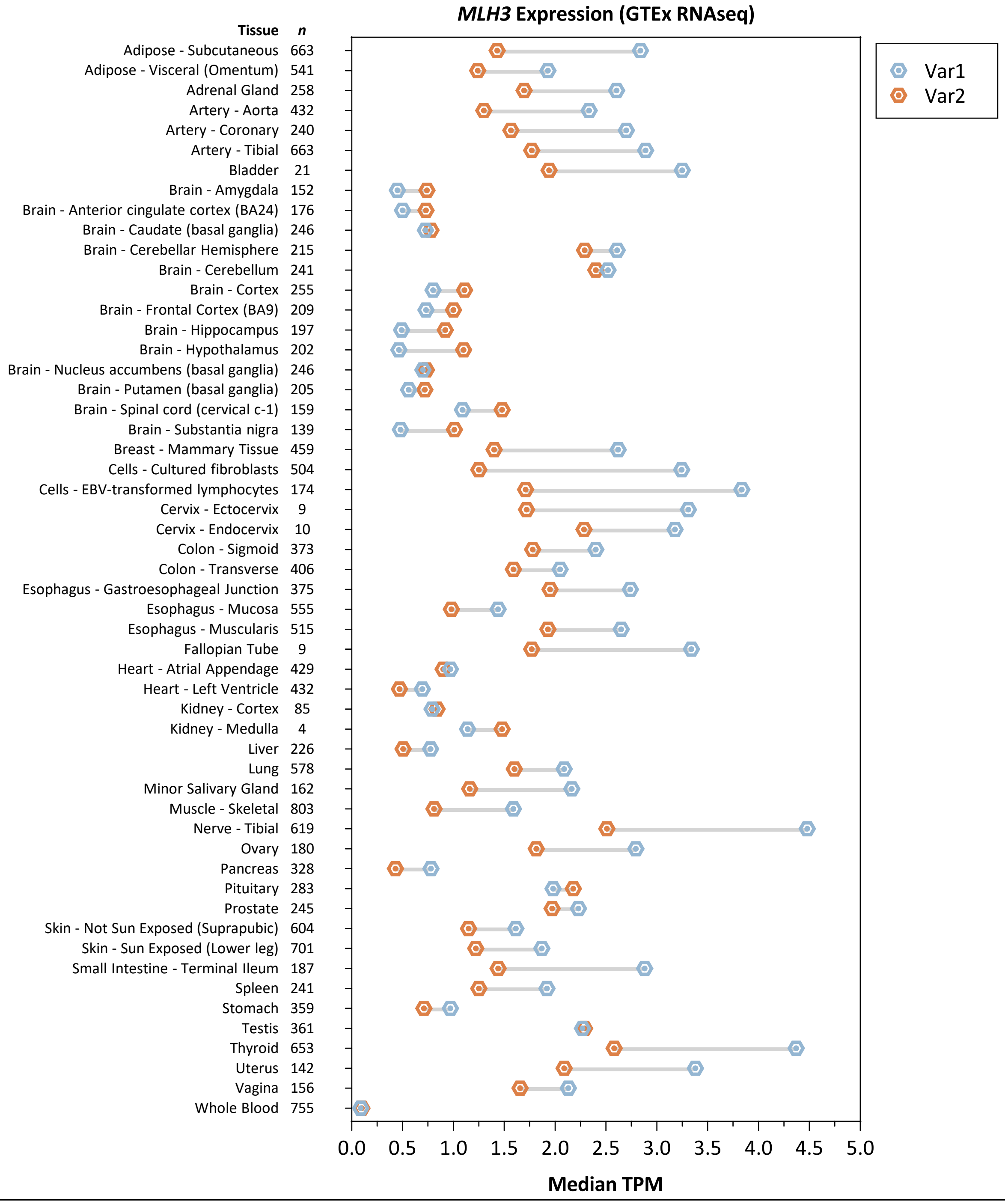


GTEx RNA-Seq data (analysis release V8, dbGaP Accession phs000424.v8.p2) was downloaded from the GTEx Portal on 07/08/2020 (GTEx_Analysis_2017-06-05_v8_RSEMv1.3.0_transcript_tpm.gct.gz). The median transcript TPM (Transcripts Per Kilobase Million) was calculated for all annotated *MLH3* transcripts (Ensembl gene version ENSG00000119684.15) across all available samples (*n* = 17,382 total samples; *n* = 4-803 samples per tissue). *MLH3* variant 1 (Var1) was calculated by combining two transcripts (ENST00000355774.6 and ENST00000556740.5), both of which, encode the full-length MLH3 protein isoform with the endonuclease domain (CCDS32123.1). *MLH3* variant 2 (Var2) represents a single transcript (ENST00000380968.6) that encodes the MLH3 protein isoform lacking the endonuclease domain (CCDS9837.1). The remaining *MLH3* transcripts detected do not represent any human consensus coding sequence (CCDS) and were excluded from this analysis.

**Figure S5. *HTT* CAG repeat size distributions in HD patient-derived fibroblasts before and after six weeks of *MLH3* SSO treatment.**

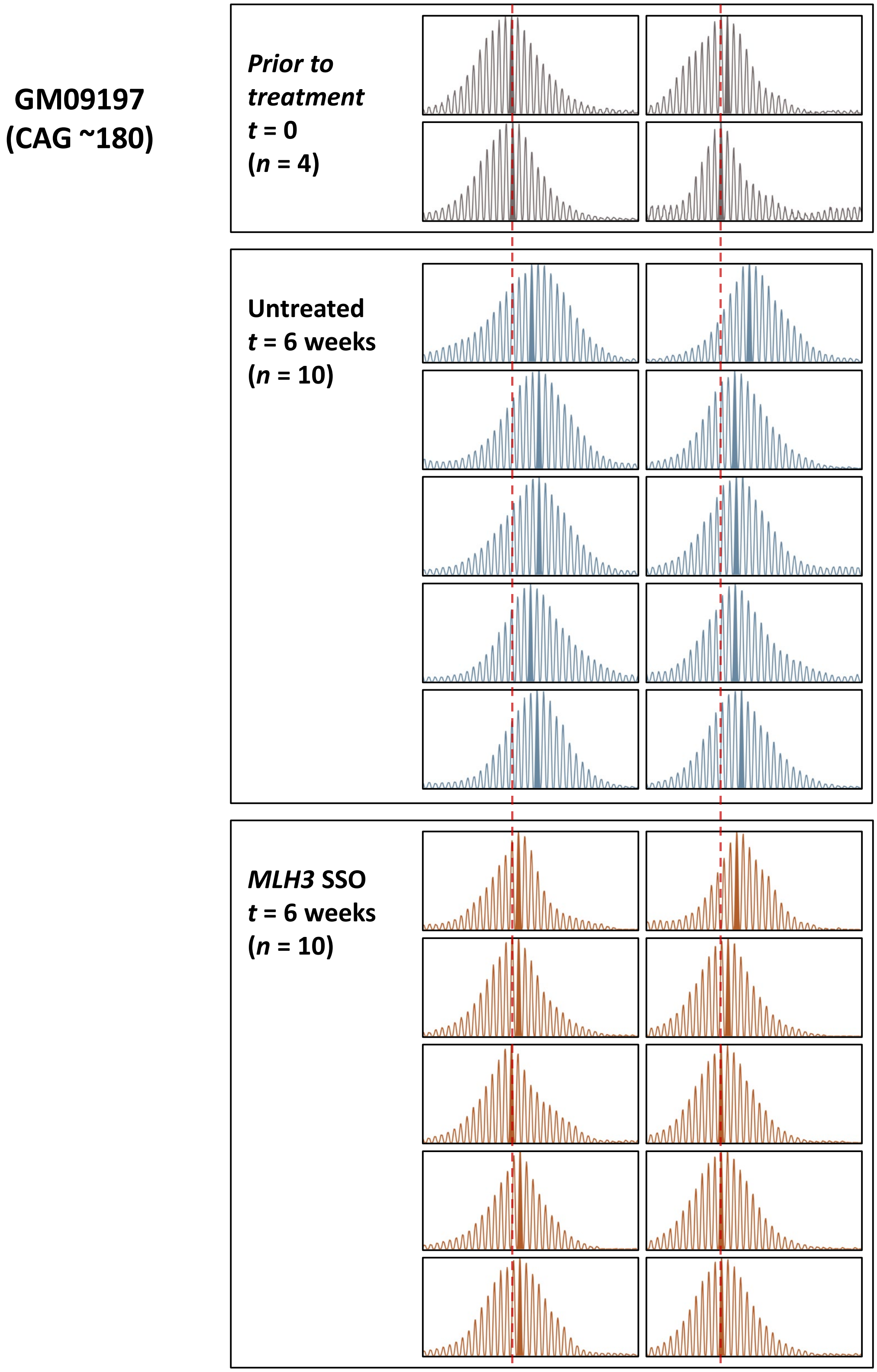


GeneMapper traces of *HTT* CAG repeat size distributions in primary HD fibroblasts (GM09197, CAG ~180) before (*t* = 0) and after six weeks of CAG expansion induction by ectopic MSH3 expression (*t* = 6weeks). During this period, cells were either treated with *MLH3* SSOs two times per week, or left untreated. Solid peaks indicate the modal CAG allele. Red lines represent the modal CAG at *t* = 0.
